# Dynamical rearrangement of human epidermal growth factor receptor 2 upon antibody binding: effects on the dimerization

**DOI:** 10.1101/752980

**Authors:** Pedro R. Magalhães, Miguel Machuqueiro, José G. Almeida, André Melo, M. Natalia D. S. Cordeiro, Sandra Cabo Verde, Zeynep H. Gümüş, Irina S. Moreira, João D. G. Correia, Rita Melo

**Affiliations:** Centro de Química e Bioquímica and Departamento de Química e Bioquímica, Faculdade de Ciências, Universidade de Lisboa, 1749-016 Lisboa, Portugal; CNC - Center for Neuroscience and Cell Biology; Rua Larga, FMUC, Polo I, 1ºandar, Universidade de Coimbra, 3004-517; Coimbra, Portugal; REQUIMTE/LAQV, Faculdade de Ciências da Universidade do Porto, Departamento de Química e Bioquímica, Rua do Campo Alegre, 4169-007 Porto, Portugal; Centro de Ciências e Tecnologias Nucleares and Departamento de Engenharia e Ciências Nucleares, Instituto Superior Técnico, Universidade de Lisboa, CTN, Estrada Nacional 10 (km 139,7), 2695-066 Bobadela LRS, Portugal; Department of Genetics and Genomics and Icahn Institute for Data Science and Genomic Technology, Icahn School of Medicine at Mount Sinai, New York, NY 10029, USA

**Keywords:** Breast cancer, dimerization inhibition, human epidermal growth factor 2 (HER2), molecular dynamics, receptor-antibody interactions

## Abstract

Human epidermal growth factor 2 (HER2) is a ligand-free tyrosine kinase receptor of the HER family that is overexpressed in some of the most aggressive tumours. Treatment of HER2+ breast cancers with the humanized monoclonal anti-HER2 antibody (Trastuzumab) revealed highly effective, encouraging the development of various HER2-specific antibodies, kinase inhibitors and dimerization inhibitors for cancer therapy. Although it is known that HER2 dimerization involves a specific region of its extracellular domain, the so-called “dimerization arm”, the mechanism of dimerization inhibition remains uncertain. However, uncovering how antibody interactions lead to inhibition of HER2 dimerization is of key importance in understanding its role in tumour progression and therapy. Herein, we employed several computational modelling techniques for a molecular-level understanding of the interactions between HER and specific anti-HER2 antibodies, namely an antigen-binding (Fab) fragment (F0178) and a single chain variable fragment from Trastuzumab (scFv). Specifically, we investigated the effects of antibody-HER2 interactions on the key residues of “dimerization arm” from molecular dynamics (MD) simulations of unbound HER (in a total of 1 µs), as well as scFv:HER2 and F0178:HER2 complexes (for a total of 2.5 µs). A deep surface analysis of HER receptor revealed that the binding of specific anti-HER2 antibodies induced conformational changes both in the interfacial residues, which was expected, and in the ECDII, in particular at the “dimerization arm”, which is critical in establishing protein-protein interface (PPI) interactions. Our results support and advance the knowledge on the already described trastuzumab effect on blocking HER2 dimerization through synergistic inhibition and/or steric hindrance. Furthermore, our approach offers a new strategy for fine-tuning target activity through allosteric ligands.

**Author summary:** Increasing insight into the genetics and molecular biology of diseases has resulted in the identification of a high number of potential molecular targets for drug discovery and development. Human Epidermal Growth Factor Receptor 2 (HER2) is one of the most relevant Epidermal Growth Factor Receptor (EGFR) members, whose overexpression has been shown to play an important role in the development and progression of certain aggressive types of breast cancer. Thus, the development of novel approaches on anti-HER2 therapies is quiet relevant. Molecular modelling and simulation can be used to bring new perspectives, both in structure-based drug design and to provide atomistic information of the intermolecular coupling dynamics between inhibitors and receptors via interactive computographic software. Considering my interest in drug design and mechanisms of action using integrated in silico approaches, herein, I have used multiple methods to evaluate how HER2 coupling to its different partners could alter this functional mechanism. The results suggest that the antibodies fragments studied show different dynamic complexes with HER2 although both could contribute to downstream of the tumour cell receptor pathways. I do believe that future research breakthroughs with aid of chemo-bioinformatics will allow a more comprehensive perception on biomedicine.

## Introduction

The human epidermal growth factor receptor (EGFR) family comprises four members: human epidermal growth factor 1 (HER1), HER2, HER3 and HER4(1). They are overexpressed in a wide variety of malignant tumours, in particular in Non-Small Cell Lung Cancer (NSCLC) (40-80 %), breast cancer (14-91 %) and head and neck cancer (80-100 %) (2). In recent years, EGFR became an important biomarker and target of anti-tumour therapy(3). Within the EGFR family, HER2 is one of the most important members and when overexpressed, represents one of the most aggressive phenotypes (approximately 20-25 % of invasive breast cancers)(4), making HER2 a well-established therapeutic target. Trastuzumab (Herceptin^®^) was the first FDA-approved antibody against HER2 and remains the gold standard for the treatment of HER2-overexpressing cancers(5). Clinical studies showed that the combination of trastuzumab with adjuvant chemotherapy substantially improves disease-free survival (DFS) and overall survival(6).

From a structural point of view, HER2 is a 1255-amino-acid transmembrane glycoprotein with 185 kDa(7, 8) known for its function as a ligand-less co-receptor. It is noteworthy that HER2 is the only member of the EGFR family that does not homodimerize under normal conditions, and its ectodomain does not directly bind ligands(9). It is activated via heterodimerization with other ligand-bound receptors, with HER3 the preferred heterodimerization partner(10) or homodimerization when it is expressed at very high levels on the cell surface(11). HER2 dimerization activates cytoplasmic kinase, which leads to autophosphorylation of its intracellular domains and thereby downstream signalling of cellular pathways. In normal cells, few HER2 molecules exist at the cell surface, and therefore few heterodimers are formed, allowing growth signals to be relatively weak and controllable. However, if HER2 is overexpressed, the dimerization level increases, resulting in enhanced responsiveness to growth factors and malignant growth(8). Structurally, HER2 comprises an extracellular domain (~600 amino acid residues, Fig 1A) with four extracellular domains (ECD) (ECDI-ECDIV), including two cysteine rich domains (ECDII and ECDIV), a transmembrane domain, an intracellular tyrosine kinase domain and a C-terminal tail with autophosphorylation sites(12).

**Fig 1.**
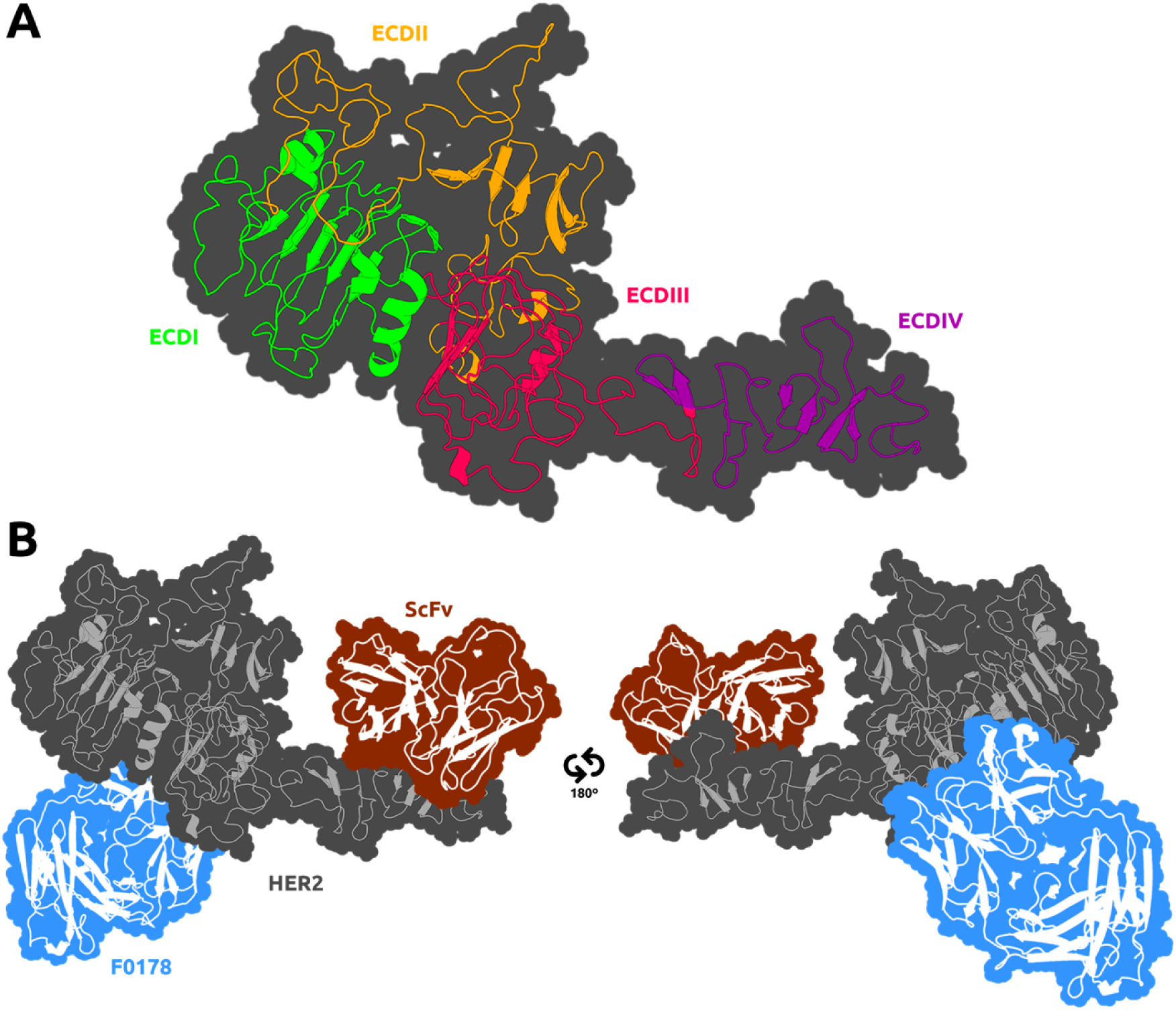
Depiction of the HER2 receptor with its four different ECD regions mapped from PDBID: 1N8Z(13) (A). The regions where both partners are located at the complex crystal structure are also shown (B). The F0178:HER2 structure was obtained from PDBID: 3WSQ(14) whereas the ScFv:HER2 structure was obtained from PDBID: 1N8Z(13). The secondary structure of the receptor and its partners is depicted as a cartoon, and the surface is shown as a contour.

The structures of the ligand-free HER2 ECDs (Fig 1A), ECDI and ECDIII, are very similar, with each containing a “β-helix” with sides, formed by three parallel β-sheets due to a high number of hydrophobic residues, mainly leucine. The spatial conformation of these two subdomains hinders the effective access in part explaining lack of ligand binding by HER2. ECDII and ECDIV also share a similar structure with small structural units held together by one or two disulphide bonds. ECDIV, a near transmembrane domain, is crucial to the stabilisation of the protein-protein interactions between HER2 and its dimerization partner(15). The structural conformation of ECDI and ECDIII brings close together, allowing ECDII projection to participate in receptor dimerization and initiation of signal transduction(16). HER2 also includes a fixed, open conformation overhand outside of the ECDII to contact, in particular, with the so-called “dimerization arm” at Δ265-288 region of its dimerization partner. This region is a short hairpin loop(17, 18) with conserved residues and plays an important role in the dimerization process(16, 18, 19). Fu *et al.* have inferred the coupling mechanism of HER2 with the antigen-binding antibody fragment (Fab) F0178 based on the three-dimensional (3D) static crystal structure of F0178:HER2(14). The authors claimed that F0178 was able to bind to HER2 through a large and highly complementary interface including both ECDI and ECDIII expressing a high heterodimerization blocking activity(14). However, the structural mechanism underlying HER2 dimerization is still unknown.

Here, we have employed a computational approach to deepen the current understanding of the binding mechanism of HER2 with specific anti-HER2 antibody fragments. For this purpose, we have selected two well-known anti-HER2-antibody fragments, namely F0178 and a single chain variable fragment (scFv) from Trastuzumab. Both antibodies inhibit HER2 with distinct mechanisms of action. The antibody fragment-HER2 complexes are depicted in Fig 1B. For the study of the ScFv:HER2 system we have considered the crystal structure of HER2 with trastuzumab Fab(13). The crystal structure showed that the trastuzumab Fab binds to the C-terminal region of ECDIV of HER2, far away from the known dimerization region (ECDII). This interaction blocks proteolytic cleavage in HER2 and indirectly affects dimerization with other HERs(20). It has been hypothesised that binding of trastuzumab could induce conformational changes providing a steric barrier avoiding kinase activation. This assembly could also promote steric blocking of HER2 by proteolytic cleavage of its extracellular region and thereby altering interaction between HER2 and other proteins(6).

*In silico* approaches are of crucial importance to clarify the structural and dynamical factors that influence protein-protein interfaces (PPIs). They allow the prediction of how these interactions occur, improving design and development of new therapeutic agents with better efficacies. Aiming at a detailed picture of the overall coupling mechanism between the different antibody fragments and the receptor as well as to understand the HER2 dimerization process, we employed computational modelling and Molecular Dynamics (MD) simulations of HER2 receptor in the apo form and upon complexation with F0178 and scFv from trastuzumab. We performed an undocumented systematic analysis of structural, energetic and dynamic characteristics, ranging from pairwise interactions formation to covariance analyses for these complexes. Moreover, we identified the most important changes occurring at these PPIs and their influence on the conformational rearrangement of the specific domains involved in dimerization, which is key to the development of new HER2-specific therapies.

## Results

### 1. Reaching equilibration in the MD simulations

To attain equilibrium in MD simulations of large protein complexes is not a trivial task. We need to evaluate the more conventional structural properties, like the root-mean-square deviation (RMSD), but we will also need to monitor larger conformational transition of the quaternary subunits that may appear at the slower timescales. The RMSD values were calculated using the Cα atoms of the receptor, with either using all ECD regions, or excluding ECDIV (Fig 2). The inclusion of the ECDIV region in the calculations greatly increased the overall RMSD value due to its high conformational flexibility as can be observed in unbound HER2 and in the F0178:HER2 complex (Fig 2A-B). However, in ScFv:HER2, the contribution of ECDIV is reduced, which was assigned to the strong interaction of the ScFv with this region (Fig 1B). Overall, the ECDI-III region equilibrates relatively fast, while the ECDIV RMSD values indicate larger movements occurring at larger timescales but of more difficult equilibration.

**Fig 2.**
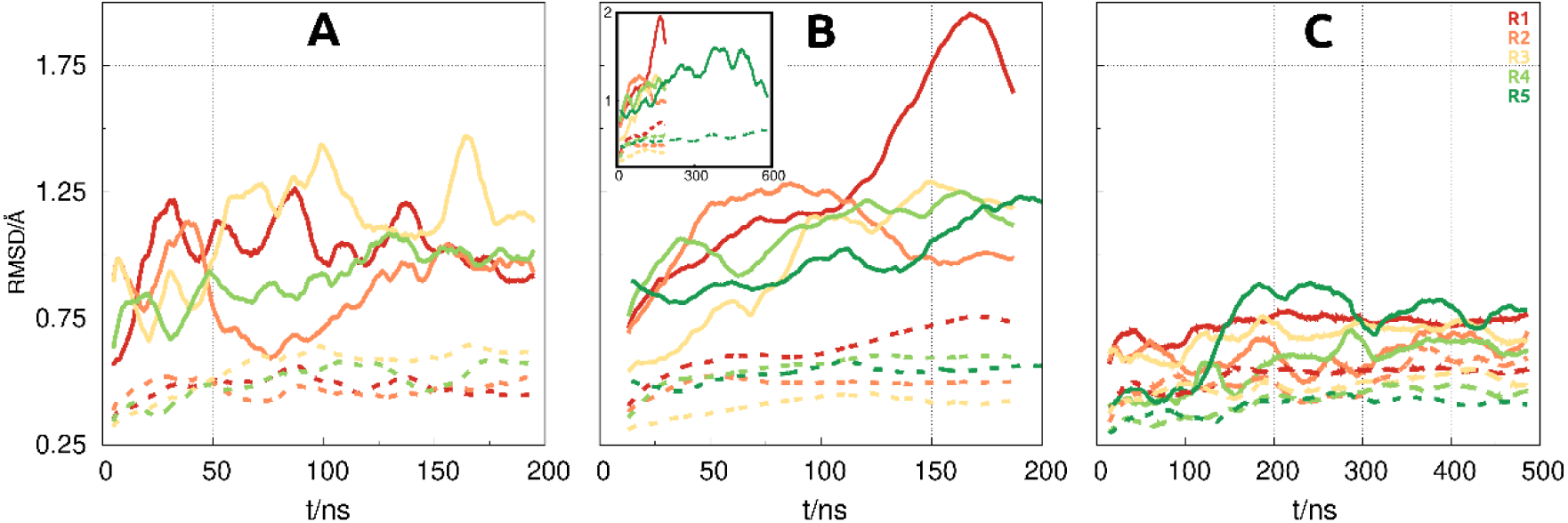
RMSD plots of the five MD simulation runs (R1-R5) for HER2 (A), F0178:HER2 (B) and ScFv:HER2 (C). Full lines depict calculations made with all four ECD regions, whereas dotted lines do not include ECDIV. The small insert in Fig B shows the full length of the longer production run (R5).

To investigate the slower conformational transitions between HER2 and its partners, we have also calculated the centre-of-mass (COM) distance between them. This distance is almost invariant in F0178:HER2 (Fig S01), since F0178 is interacting directly with the ECDI-ECDIII region of HER2 (Fig 1B), inducing no significant domain movement. In the ScFv:HER2 system (Fig S02), we observed that the ScFv, upon binding, induced an opening/closing movement of the ECDIV towards the ECDI-III domains. As described in the methods section, this movement, which varies in the hundreds of nanoseconds time scale, was the reason why we have extended the MD simulations to improve its sampling.

Brought together these results showed that ScFv position varies between two extremes, depending on receptor proximity (Fig 3). The maximum COM distance leads to an opening of the cleft between ScFv and the HER2 ECDI-III domains (Fig 3A), an intermediate position, which is similar to the initial conformation (Fig 3B); and a minimum distance, which leads to cleft closure and a direct interaction of ScFv with ECDI-III (Fig 3C). Next, we selected the equilibrated regions of the MD simulations based on the RMSD and COM distances data. In the HER2 control and the F0178:HER2 complex simulations, these equilibrated rather fast (30-50 ns), whereas in the ScFv:HER2 system, due to the large domain movements, some replicates took up to 150 ns to reach equilibrium. In this system, all replicates seemed to converge to a different COM distance, indicating that the use of multiple replicates was crucial to improve the sampling of this movement.

**Fig 3.**
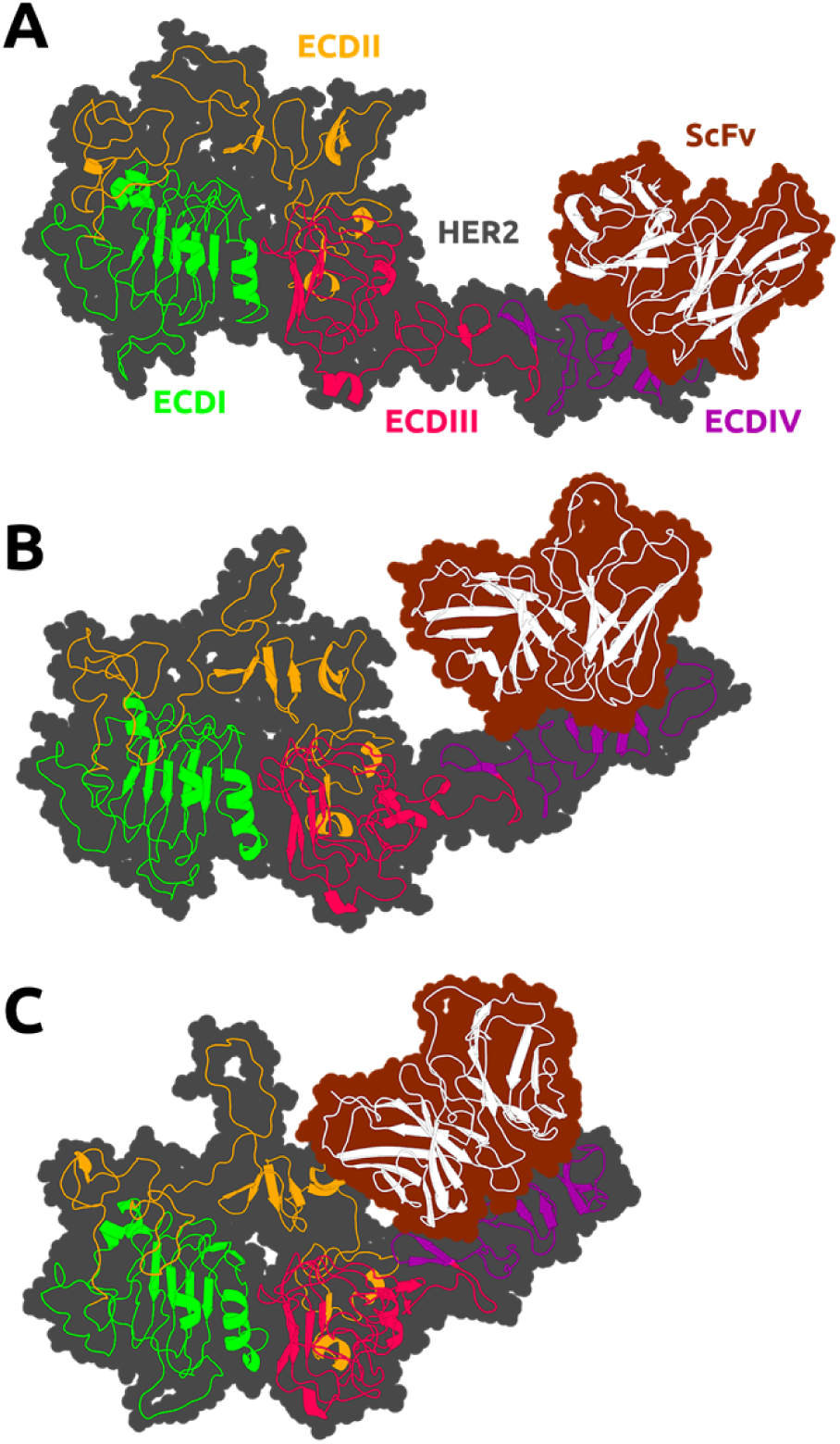
Snapshots of the ScFv:HER2 system in open (A), intermediate (B) and closed (C) conformations. The secondary structure of the receptor and its partners is depicted as a cartoon, and the surface is shown as a contour.

### 2. HER2 Conformational reorganisation

After the equilibration evaluation of the MD simulations, we identified the equilibrium conformational space and performed a deeper analysis of HER2 conformation reorganisation after antibody binding. Briefly, it has been previously described that HER2-ECDII is the main domain involved in the dimerization process(19, 21). Therefore, we have used an integrated approach of multiple methods to evaluate how HER2 coupling to its different partners could alter this functional mechanism. In addition, we have performed cross-correlation analysis for both complexes (Fig 4).

**Fig 4.**
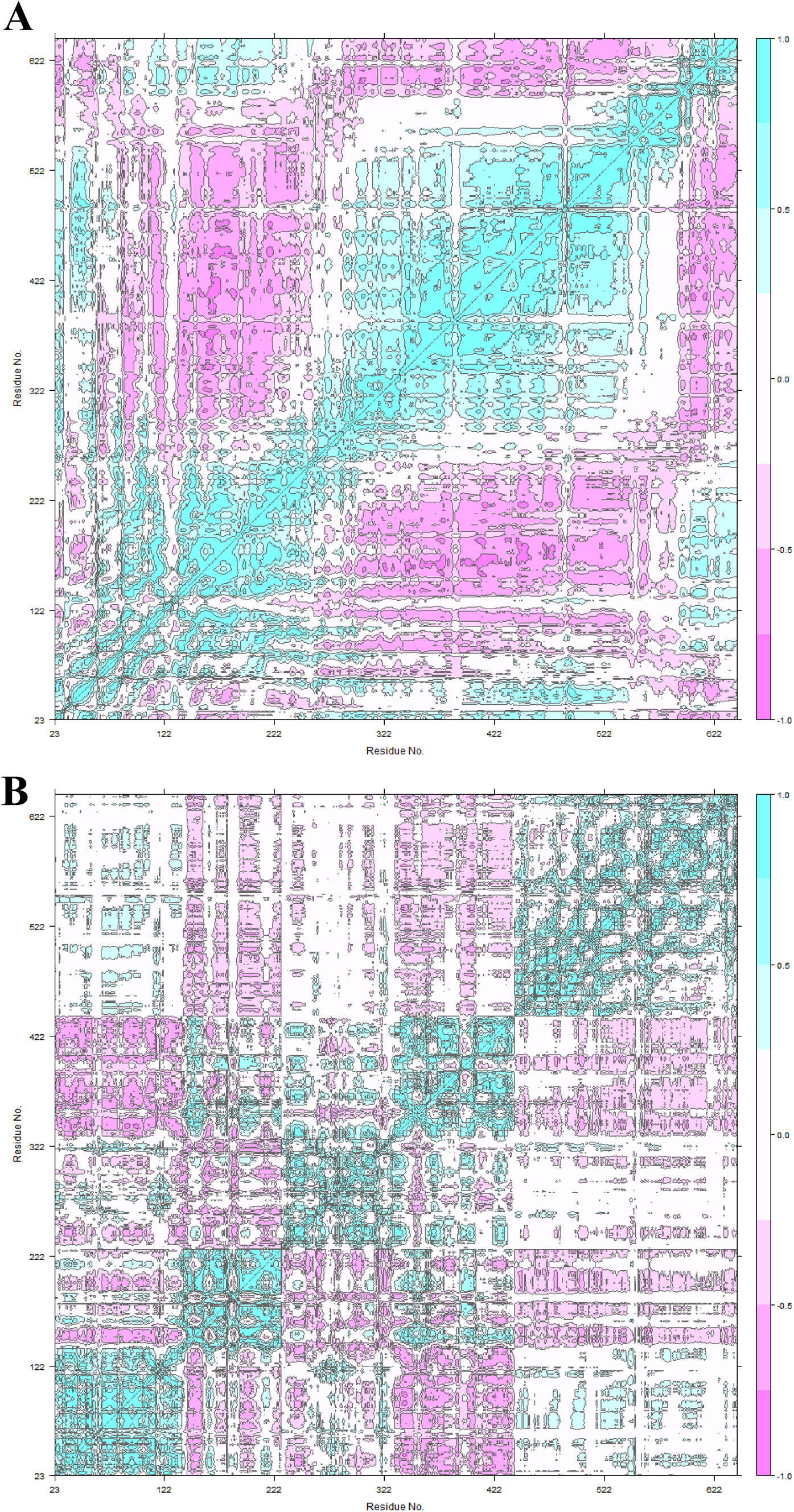
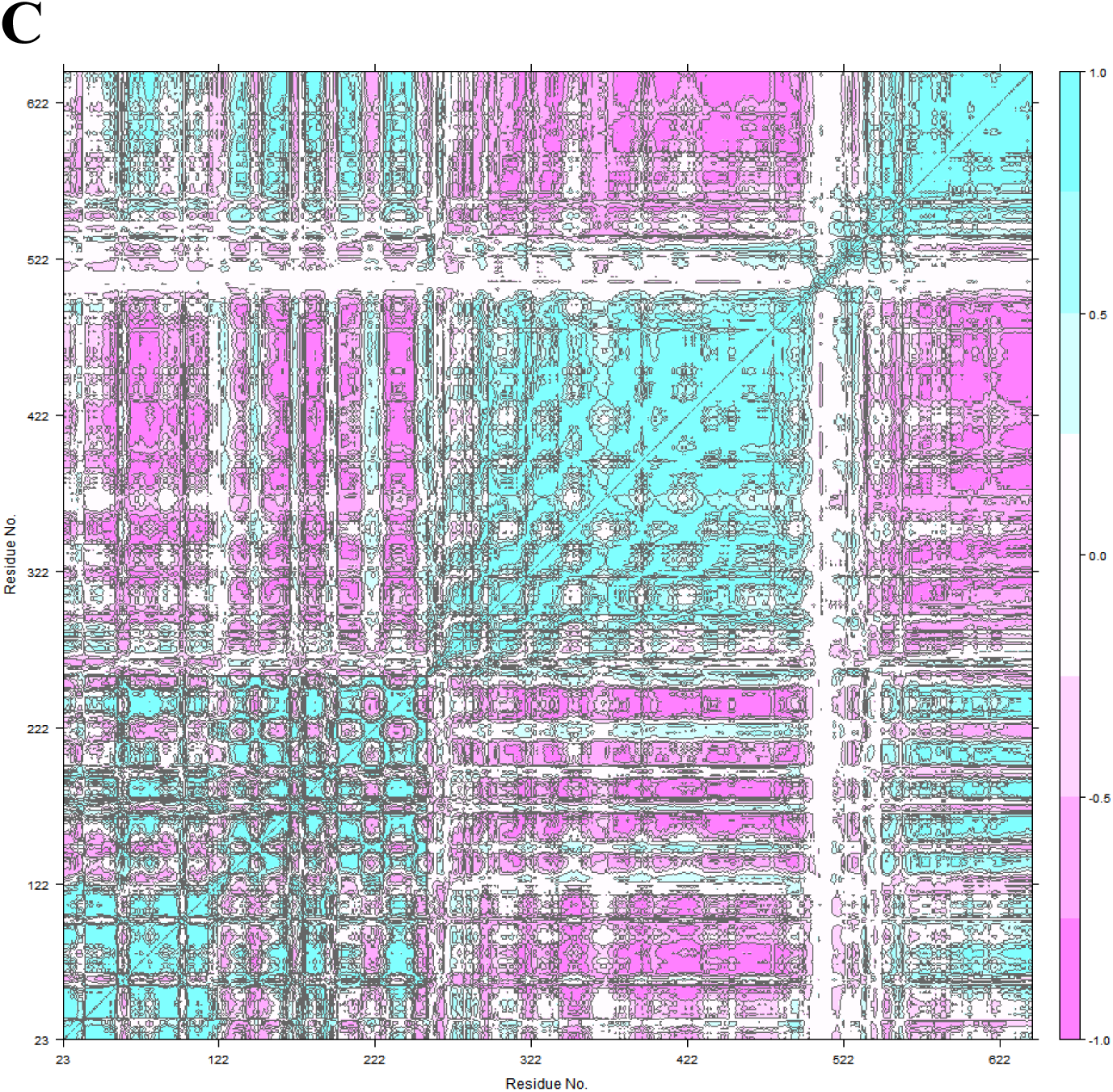
Cross-correlation Analysis. A: unbound HER2; B: HER2 at F0178:HER2 and C: HER2 at scFv:HER2.

For the unbound HER2 (Fig 4A) we found a mixed correlation spectrum (both positive and negative) for the 23-186 region (ECDI) and negative correlation values (excluding ECDIV) for the rest of the protein. Fu *et. al.* identified HER2 sub-region ASN498-ASP502 (ECDIII) as particularly altered due to F0178 binding even though it is positioned slightly apart(14). Cross-correlational analysis for F0178:HER2 shows strong and weak, respectively, negative correlation when comparing the 23-186 region (ECDI, especially sub-region 23-100) with ECDII (187-363) and ECDIII (364-530) regions. The negative correlation of ECDIII with ECDI (1-186 region) and ECDIV (531-626 region) underscores the importance of ECDIII on the F0178:HER2 interaction. Another very interesting aspect is the positive correlation between ECDIII and ECDII (187-363 region).

Concerning ScFv:HER2 (Fig 4C), a strong positive correlation between ECDIV tail (above 800) and ECDI and ECDII revealed the importance of ECDIV on the ScFv:HER2 interaction. The strong negative correlation of ECDII and ECDIII, as opposed to HER2 movement when is unbounded (Fig 4A) suggested a conformational change with scFv binding at ECDIV. This behaviour is in line with that shown in Fig 3 in which ScFv:HER2 complex varies its position between two extremes.

To extract the relevant information from the conformational rearrangement observed during the MD simulations, we performed a PCA analysis for the F0178:HER2 and ScFv:HER2 systems. These calculations were performed using all replicas and considering only the equilibrated regions of the MD simulations. Conformational clustering was also used to identify and further illustrate the most populated conformers sampled in the MD simulations. For both systems, over 78-95 % of the movement was described by the first 3 principal components, which allowed us to positively identify the most relevant conformational transitions (Fig 5)

**Fig 5.**
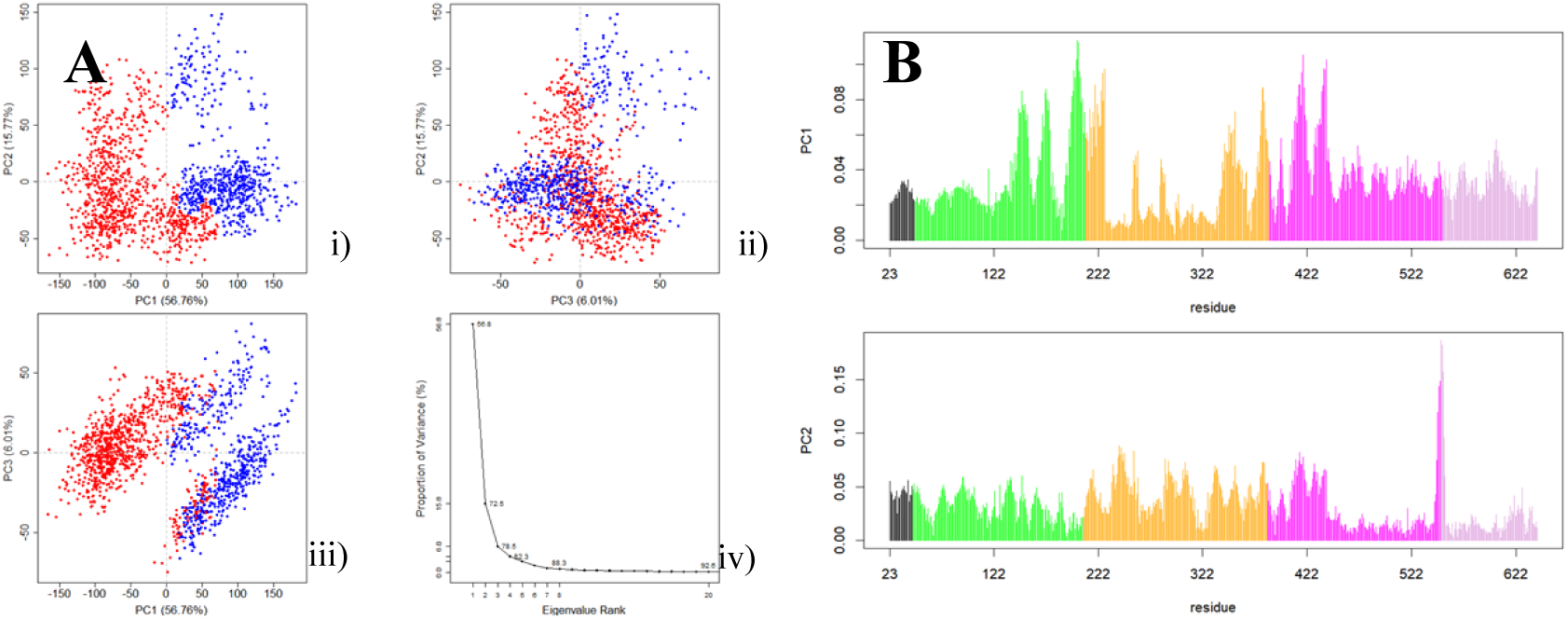
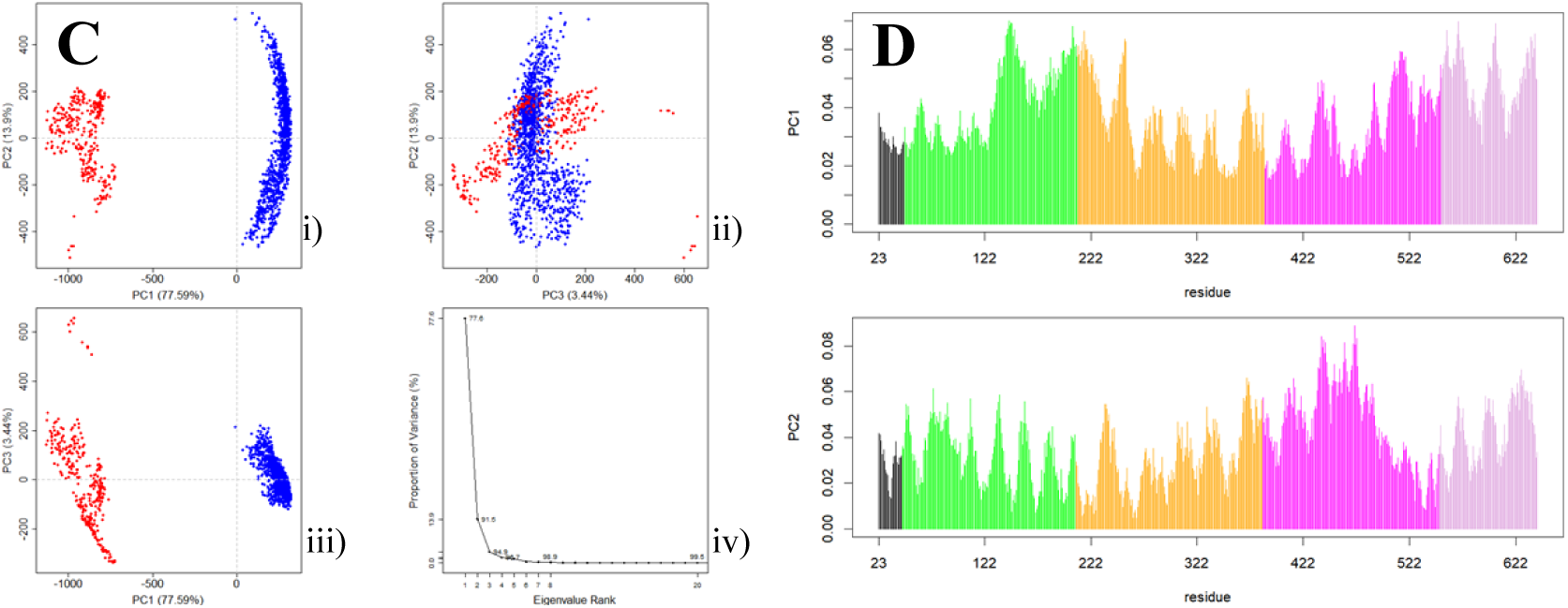
PCA clustering results of the MD simulation trajectories of the complexes. i. Principal components (PC) 1 and 2. ii. PC1 and PC3. iii. PC2 and PC3. iv. Percentage of variance explained by the first 10 PC: (A) F0178:HER2, (C) scFv:HER2. Contribution of each residue to the first two principal components: (B) F0178:HER2, (D) scFv:HER2.

The PCA analysis for the F0178: HER2 MD simulations excluded the ECDIV tail, since its large movement (Fig 2B) would mask any other movement by the other domains. Assuming this approach allowed us to get insights about conformational details in regions otherwise inaccessible. Fig 5A shows slightly distinct conformational states mainly dependent on the ECDII at 187-363 region. Remarkably, the ECDII (187-363 region) residues had a significant contribution to the first two principal components (Fig 5B). We also calculated the conservation scores using the ConSurf web-server(22). Fig S03 shows a high conservation score for GLU352, VAL353 and ARG354 at F0178:HER2 complexes, which is in accordance with the higher contribution of ECDII region for PC1.

PCA analyses suggested two main conformational states for the scFv: HER2 MD simulations (Fig 5C). In particular, the contribution of the ECDII residues to PC1 and PC2 (Fig 5D) suggested that the dimerization region was indeed affected by binding. Fig S03 also supports this behaviour with a high rate of the conservation pattern at ECDII, namely at dimerization arm (GLU265-ARG288). Paradoxically, the role of the ECDII-HER2 residues in the interface-interaction network of both complex systems were not considered in the literature (13, 14). To further aid interpretation, we produced a trajectory that interpolates between the most dissimilar structures in the distribution along PC1 for both systems (Fig S04). The movement of the “dimerization arm” at ECDII is rather evident, as well as at ECDI. The role of this region for HER2 coupling mechanism has not been previously described although the blocking ability of HER2 heterodimerization is well recognised(14). We propose that the HER2 ECDII conformational change is coupled with this allosteric regulation of F0178:HER2.

Concerning the scFv:HER2 complex system, we observed an approximation between ECDII and scFv compared to the initial 3D crystallographic structure (PDBID 1NZ8(13)) (Fig S04). This is an important biological feature since it is well-known that the ‘heart-shaped’ domain II open conformation is essential for dimerization^17^.

### 3. The impact of the ScFv and F0178 on the conformational space of HER2

Based on the analysis of the conformational changes that occurred during the MD simulations of the systems, the ScFv and F0178 antibody fragments seem to induce conformational changes in HER2, which might alter its dimerization propensity. The most impactful changes are those that alter the receptor surface, in particular, in the regions known to be involved in dimerization. These surface changes are inevitably coupled with variations in the solvent exposure of certain residues. Therefore, we calculated the solvent accessible surface area (SASA) of each HER2 residue in all systems to calculate significant differences between complexes and the unbound receptor. The residues with significant differences in SASA (difference > 5 % and standard error is smaller than the difference value) due to the presence of a partner are associated with a direct interaction and should correlate well with the interfacial region (Fig 6). As expected, the residues with the largest shift in SASA in F0178:HER2 are located in ECDI, ECDII and ECDIII (Fig 1B and Fig 6A), whereas in ScFv:HER2 these residues are mainly located in ECDII and ECDIV (Fig 1B and Fig 6B). In the case of ScFV:HER2, the direct interaction of the ligand with ECDII is due to the cleft closure (Fig 3C), observed in several simulations (Fig S02). The large error bars in F0178:HER2 were attributed to insufficient sampling and/or an increase variability in the interfacial region of the complex.

**Fig 6.**
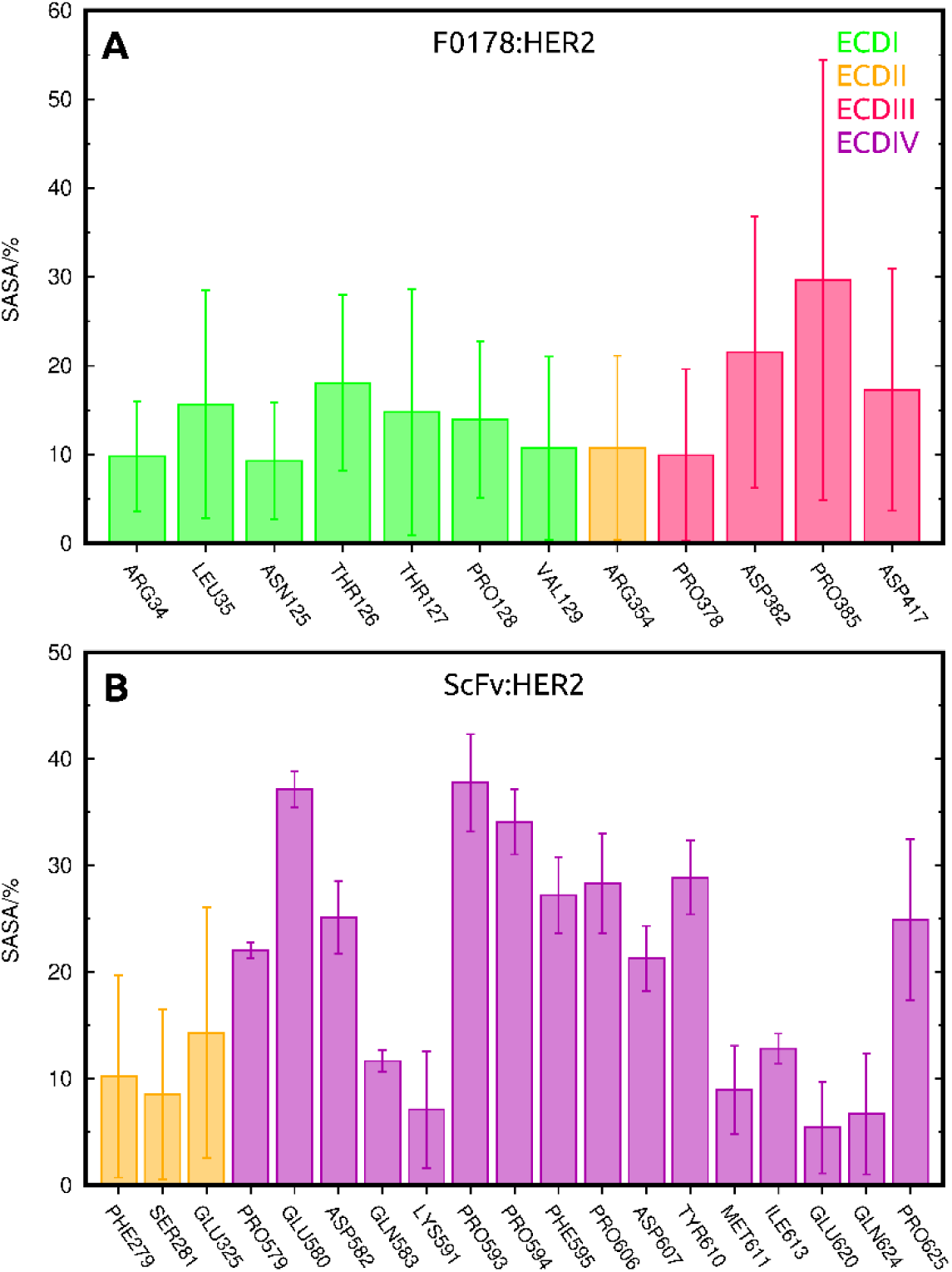
Residues with significant differences in SASA due to the presence of a partner, coloured by their respective ECD in F0178:HER2 (A) and ScFv:HER2 (B).

We also identified several residues whose SASA changed between unbound HER2 and its complexes through an indirect, or allosteric, effect of the antibody fragment. This was attained by calculating the SASA differences between F0178:HER2/ScFv:HER2 and HER2 trajectories, ignoring the physical presence of the ligands. The residues identified with this procedure (Fig 7 and Fig S05) were different in both complexes, but located predominantly in ECDII, including the so-called “dimerization arm” region (TYR274, GLU280 and MET282 in F0178:HER2; and CYS268 in ScFv:HER2).

**Fig 7.**
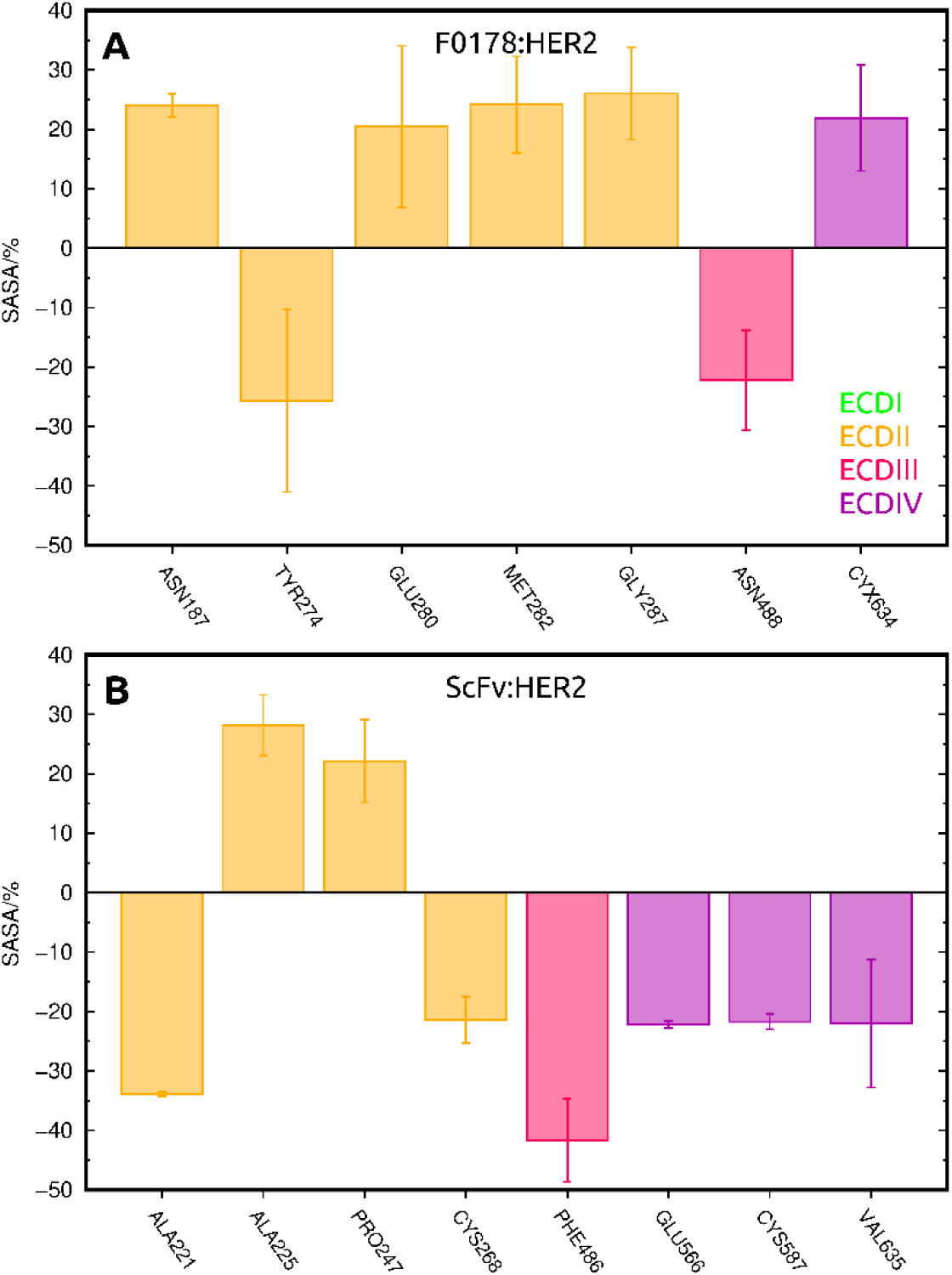
Residues with the largest difference in SASA due to allosteric effects, colored by their respective ECD in F0178:HER2 (top) and ScFv:HER2 (bottom).

Our results indicate that F0178 and scFv form fairly distinct dynamic complexes with HER2. The binding of F0178 seems to provoke a marked allosteric effect on the “dimerization arm” residues at ECDII, which is the dimerization region of HER2(16). A similar allosteric effect could be observed for scFv, which was complemented with a direct interaction with the ECDII region. This interaction map obtained from our simulations may help explain the strong effect of scFv on HER2 homodimerization(6), opening new therapeutic avenues for targeting HER2 inhibition.

## Discussion and Conclusions

HER2 plays a key role in the HER family due to its specificity to interact with other HER receptors through a complex signalling network to regulate cell growth, differentiation and survival. HER2 overexpression occurs in approximately 15–30 % of breast cancers and 10–30 % of gastric/gastroesophageal cancers and serves as a prognostic and predictive biomarker. The easy accessibility to the four HER2 extracellular domains (ECD) turns this receptor into a highly relevant target for the selective delivery of anti-tumour drugs as well as imaging agents. ECDI and ECDIII can form a binding site for potential HER2 ligands, whereas the cysteine-rich ECDII and ECDIV are known to be involved in the homodimerization and heterodimerization processes. Furthermore, ECDII, in particular the “dimerization arm” region (Δ265-288), is believed to be the main contributor to dimerization. Although crystal structures are known, the dynamics of the binding of different antibodies to this receptor has remained unclear. In this work, we have applied an integrated framework of molecular modelling tools and techniques as well as MD simulations analyses to address important structural factors that provide atomistic information of the intermolecular coupling dynamics between HER2 and two different HER2 inhibitors, namely F0178 Fab and scFv from trastuzumab. Both are well-studied anti-HER2 antibody fragments that act at different ECD domains of HER2. F0178 interacts with both ECDI and ECDIII domains, while scFv binds at ECDIV domain. Concerning the ScFv:HER2 complex, the highest contribution to the PPI stability occurs in the terminal part of ECDIV.

In addition to the detailed residue and pairwise interaction analyses of HER2-antibody complexes, an important contribution of our work is on the conformational changes that occur upon binding. Our simulations suggested that F0178 and scFv form distinct dynamic complexes with HER2. The binding of F0178 seems to induce a marked allosteric effect on the “dimerization arm”, whereas scFv binding leads to major conformational changes on HER2 structure (ECDI approaches ECDII and ECDIII). We observed a drastic movement of ECDII towards scFv, establishing a new possible interface capable of meaningful and important interactions with the antibody. This newly discovered conformational organisation of HER2 upon scFv coupling could indeed explain its role on dimerization as suggested by Hudis *et al.*(6) and herein we give a pioneer view of this crucial dynamical process. The integrated analysis framework applied here has unveiled the role of F0178 and scFv from trastuzumab in HER2 dimerization. Our results provide a new perspective for future approaches to the selection of effective potential drugs, since the conformational changes triggered by the ligands can have positive effects downstream of the tumour cell receptor pathways.

## Experimental section

### 1. Model Construction

The 3D structure of HER2 extracellular domain was constructed by homology modelling using the MODELLER package(23), and the protein template NP_004439.2 (PDBID: 3N85A)(24) with the sequence 301598662. The loop regions LEU99:SER133; THR245:CYS264 and ASP121:GLY136 were further refined with ModLoop(25, 26). The complexes of HER2 with antibody fragments were retrieved from the correspondent PDB files, specifically F0178:HER2 from PDBID: 3WSQ(14) and ScFv:HER2 from trastuzumab with PDBID: 1N8Z(13). The protonation states of residues in the physiological pH range were determined with the PROPKA methodology (27–29), which computes p*K*_a_ values of ionisable residues by accounting for the effect of the protein environment. PKA2PQR(27) was applied to convert PDB files in PQR files by AMBER ff99 force field(30). All histidine residues were considered deprotonated at physiological pH conditions save for HIS 349, which was assigned as positively charged.

### 2. Molecular Dynamics Simulations

MD simulations of HER2, F0178:HER2 and ScFv:HER2 were performed using GROMACS 2018.3(31–33) and the Amber ff99SB-ILDN force field (34). Each complex was solvated by a TIP3P water box with 12 Å buffering distance to the boundary, which was verified throughout the simulations. The simulation systems were kept neutral by adding the necessary counterions. Simulations were performed in the *NPT* (isothermal-isobaric) ensemble. Temperature coupling was performed using the v-rescale thermostat(35) at 300 K, with a coupling constant of 0.1 ps, while an isotropic Parrinello-Rahman barostat(36, 37) was used to keep the pressure constant at 1 bar using a coupling constant of 2.0 ps and a compressibility of 4.5 × 10^−5^ bar^−1^. Long range electrostatic interactions were computed using the particle mesh Ewald (PME) method (38, 39) with a Fourier grid spacing of 0.16 nm and a cut-off of 1.0 nm for direct contributions. Lennard-Jones interactions were computed using a nonbonded neighbour pair list with a cut-off of 1.0 nm, enabling the use of the Verlet scheme (40). Solute bonds were constrained using the Parallel LINear Constraint Solver, P-LINCS (41), and solvent molecules were constrained using SETTLE algorithm(42). The initial system energy was first minimised for 50.000 steps, followed by two equilibrations of 250 ps. The HER2 system was then initialised for 10 ps using position restraints on all protein heavy atoms with a force constant of 1000 kJ nm^−2^ mol^−1^, while the remaining systems were initialised for 25 ps without restraints.

Initially, four production runs were performed for each system, each 200 ns-long. In addition, for the F0178:HER2 and ScFv:HER2 complexes, a longer production run was also performed (~500 ns-long). However, when analysing the centre-of-mass (COM) distance between the partners in the HER2:ScFv system (Fig S02), we decided to extend the length of all production runs to 500 ns, in order to obtain better sampling of this motion.

The equilibrated regions for each system (Table S03, in the Supporting Information) was replicate/system dependent, and the decision was based on the RMSD and COM distance variations. The RMSD calculations were performed using the Cα atoms either in all extracellular domains, or in the ECDI-ECDIII region, as shown in Fig 2. The COM distance plots (Fig 3 and Figs S01-02) were obtained by calculating the centre-of-mass of the HER2 and either ScFv or F0178, using all atoms in the receptor and each of its partners.

### 3. Solvent-accessible surface area analysis

We calculated the solvent-accessible surface area (SASA) of each individual residue in HER2, ScFv:HER2 and F0178:HER2 using the equilibrated regions of each replicate. In the unbound HER2, for instance, this resulted in four average values of SASA per residue, which were then used to calculate the mean and standard error of the mean (SEM) using a jackknife technique. We then converted these absolute SASA values into percentages by calculating the maximum value a fully exposed residue of each type would have. With these percentages, we excluded all residues whose solvent-access was below 20 %, as they were considered mainly internalised.

#### 3.1 SASA analysis of interfacial residues

The identification of residues in the interface between HER2 and each of its partners was performed by calculating the SASA of each residue of the receptor in the complex, with and without its partner (e.g., the residue coloured in yellow depicted in Fig S06). When comparing the SASA values of each complex in the presence and absence of the antibody, we were able to exclude all residues whose SASA was not altered, *i.e.* residues which are not interacting directly with the partner. Finally, we only reported residues whose difference in SASA was above 5 % and whose SEM was lower than this difference.

#### 3.2 SASA analysis of allosterically modulated residues

To identify allosterically modulated residues, we compared the SASA of all residues in each complex with the same residue in unbound HER2. This SASA difference identified which residues became more exposed or occluded by an indirect effect of the antibody (*e.g.* the residue coloured in green depicted in Fig S06). As in the previous protocol, residues whose difference in SASA between systems was below 5 % and whose SEM was higher than the difference itself were excluded.

### 4. Principal Component Analysis

Principal Component Analysis (PCA)(43) is a useful statistical technique able to reveal the more important patterns in a dataset. The most relevant information from a dataset are extracted as eigenvalues or principal components (PCs). Internal correlation motions between residues of interface complexes were investigated by performing a cross-correlations analysis on the MD trajectory given a covariance matrix in Cartesian coordinate space by Eq. (1).

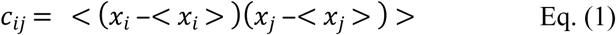

Where *x*_*i*_ and *x*_*j*_ are coordinates of the *i*^th^ and *j*^th^ atoms in the system and <…> denotes trajectory average(44). The matrix *c*_*ij*_ symmetric can be diagnosed by using orthogonal coordinate transformation matrix *T*, transforming the matrix *c*_*ij*_ into a diagonal matrix Λ of eigenvalues λ_*i*_ by Eq. (2):

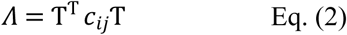

Where each eigenvector of the matrix gives the direction, along which a concerted motion arises, and the eigenvalue gives the magnitude of fluctuations along this direction. The PCA calculations were performed using bio3d(45).

### 5. Evolutionary Conservation of Interfacial Residues

Sequence conservation was retrieved for all interfacial residues in HER2 using Consurf (46), which is based on the Rate4Site algorithm(47). MAFFT(48, 49) was used for multiple sequence alignment (MSA), using the BLAST(50) on the UNIREF90 database(51). Each MSA comprised at most 150 sequences, with homology values ranging from 35 % to 95 %.

## Funding

I. S. Moreira acknowledges support by the Fundação para a Ciência e a Tecnologia (FCT) Investigator programme - IF/00578/2014 (co-financed by European Social Fund and Programa Operacional Potencial Humano). This work was also financed by the European Regional Development Fund (ERDF), through the Centro 2020 Regional Operational Programme under project CENTRO-01-0145-FEDER-000008: BrainHealth 2020, and through the COMPETE 2020 - Operational Programme for Competitiveness and Internationalisation and Portuguese national funds via FCT, under project POCI-01-0145-FEDER-007440. R. Melo acknowledges support from the FCT (SFRH/BPD/97650/2013). R.Melo, S. Cabo Verde and J. D. G. Correia gratefully acknowledge support from the FCT through projects UID/Multi/04349/2019, PTDC/QUI-NUC/30147/2017 and PTDC/QUI-OUT/32243/2017. A. Melo and M. N. D. S. Cordeiro acknowledge support from FCT/MCTES by project UID/QUI/50006/2019 (LAQV@REQUIMTE). Z. H. Gümüş acknowledges financial support from LUNGevity Foundation and start-up funds from the Icahn School of Medicine at Mount Sinai. Z. H. Gümüş, I. S. Moreira and R. Melo also acknowledge support for computational resources and staff expertise provided by Scientific Computing at the Icahn School of Medicine at Mount Sinai. M. Machuqueiro acknowledges support from FCT through contract grant CEECIND/02300/2017 and projects UID/MULTI/04046/2019, UID/MULTI/00612/2019, and PTDC/BIA-BFS/28419/2017.

## Supporting information

**Fig S01 - Time evolution of the COM distance between HER2 and F0178.** The smaller plot shows the full length of the larger production run for this system. Data was smoothed using a floating average window of 10 ns.

**Fig S02** - **Time evolution of the COM distance between HER2 and ScFv.** Data was smoothed using a floating average window of 10 ns.

**Fig S03** - **Normalised conservation scores of interfacial residues from ECDII-HER2 when coupled to F0178 (•) and scFv (+) antibody fragments.** The light blue bars highlight the so-called “dimerization arm” residues. The light blue and dark blue dashed lines represent the average conservation score for dimerization arm and non-dimerization arm residues, respectively.

**Fig S04. Interpolated structures along PC1: (A) F0178:HER2, (B) scFv:HER2.** Colour scale from blue to red depict low to high atomic displacements. The dashed box highlights the “dimerization arm” region at ECDII.

**Fig S05** - **Location of the residues that showed a significant shift in SASA due to allosteric effects.** The regions coloured in red show the residues in HER2 whose SASA was affected by the presence of ScFv, while the regions in blue show the residues in HER2 whose SASA was affected by F0178. Also shown are the initial positions of both partners (semi-transparent). The image on the right is the mirror-image of the one on the left. The secondary structure of the receptor and its partners is depicted as a cartoon, and the surface is also shown as a contour.

**Fig S06** - **Simplified depiction of the SASA analysis.** The dotted lines are a simplified depiction of the region used in the SASA calculation. Two specific residues coloured in green and yellow are also included. The residue in yellow is located in the interface region between HER2 and ScFv and should have a noticeable difference in SASA if ScFv is included or excluded from the calculation in the complex trajectory. The residue in green is located in a region where the presence of ScFv could only be felt indirectly when performing differences between the bound and unbound trajectories

**Table S01 - MD equilibrated regions for each system and replicate in nanoseconds.**

